# Assessing the ecological niche and invasion potential of the Asian giant hornet

**DOI:** 10.1101/2020.05.25.115311

**Authors:** Gengping Zhu, Javier Gutierrez Illan, Chris Looney, David W. Crowder

**Author notes:** Author Contributions All authors designed the study and wrote the manuscript, GZ conducted the analyses. **Competing Interest Statement:** None.

## Abstract

The Asian giant hornet (*Vespa mandarinia*) is the world’s largest hornet. It is native to East Asia, but was recently detected in British Columbia, Canada, and Washington State, USA. *Vespa mandarinia* are an invasion concern due to their potential to negatively affect honey bees and act as a human nuisance pest. Here, we assessed effects of bioclimatic variables on *V. mandarinia* and used ensemble forecasts to predict habitat suitability for this pest globally. We also simulated potential dispersal of *V. mandarinia* in western North America. We show that *V. mandarinia* are most likely to invade areas with warm to cool annual mean temperature but high precipitation, and could be particularly problematic in regions with these conditions and high levels of human activity. We identified regions with suitable habitat on all six continents except Antarctica. The realized niche of introduced populations in the USA and Canada was small compared to native populations, implying high potential for invasive spread into new regions. Dispersal simulations showed that without containment, *V. mandarinia* could rapidly spread into southern Washington and Oregon, USA and northward through British Columbia, Canada. Given its potential negative impacts, and the capacity for spread within northwestern North America and worldwide, strong mitigation efforts are needed to prevent further spread of *V. mandarinia*.

## Introduction

The Asian giant hornet (*Vespa mandarinia* Smith) is the world’s largest hornet and is native to temperate and sub-tropical Eastern Asia (Fig. 1A), where it is a predator of honey bees and other insects (Matsuura & Sakagami 1973; Archer 1995; McGlenaghan et al. 2019). Coordinated attacks by *V. mandarinia* on beehives involve pheromone marking to recruit other hornets, followed by rapid killing of worker bees until the hive is destroyed (McClenaghan et al. 2019). Japanese honey bees (*Apis cerana*) co-evolved with *V. mandarinia* and have defensive behaviors to counter these attacks, including recognizing and responding to marking pheromones and “bee ball” attacks on hornet workers (Sugahara et al. 2012; McGlenaghan et al. 2019). *Apis mellifera*, the European honey bee, however, has no co-evolutionary history with *V. mandarinia* and lacks effective defensive behaviors, making it highly susceptible to attack (McGlenaghan et al. 2019).

**Figure 1.**
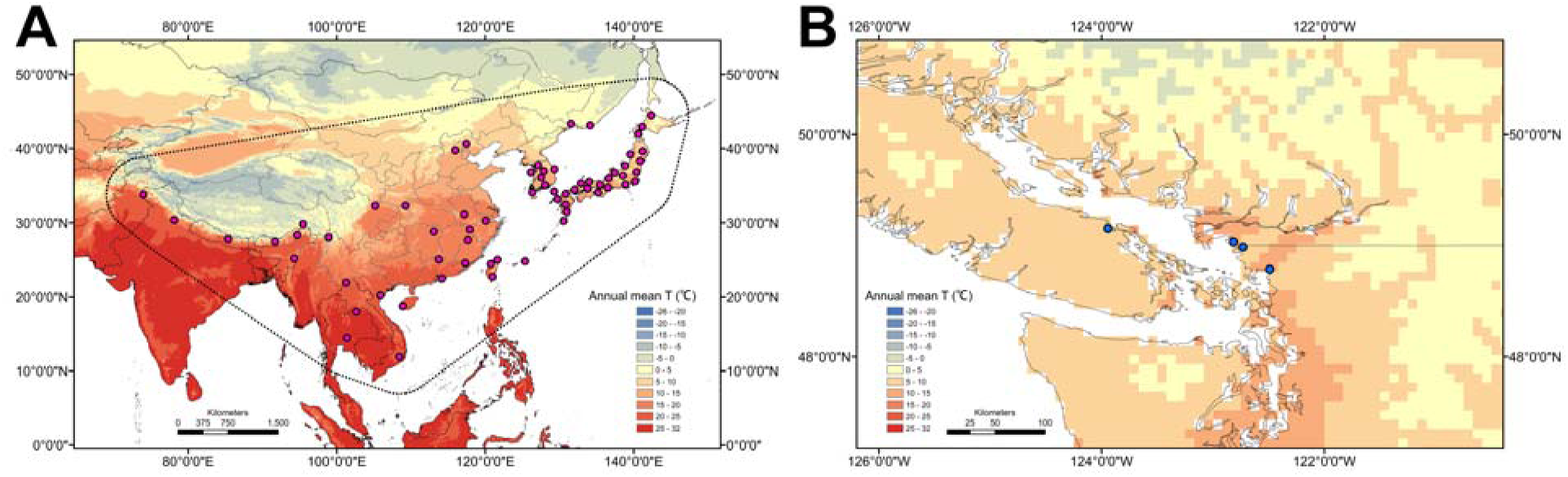
Present distribution of Asian giant hornet in (A) native and (B) introduced regions. In (A) points denote trimmed records used to fit models.

In September 2019, a nest of *V. mandarinia* was found on Vancouver Island in Canada, and two workers were found 90 km away in Washington State, USA, later that year (USDA 2019) (Fig. 1B). The introduction of Asian giant hornet into western North America is concerning because of the vulnerability of *A. mellifera*, which is widely used for crop pollination, to hornet attacks. Predation by *V. mandarinia* on *A. mellifera* in Asia causes major losses (McGlenaghan et al. 2019). *Vespa mandarinia* is also medically significant, and can deliver painful stings and large doses of cytolytic venom. Multiple stings can be fatal even in non-allergic individuals, although recent mortality rates are much lower than the historic reports of more than 30 yearly deaths in Japan (Yanagawa et al. 2007). Currently, *V. mandarinia* is listed as a quarantine pest of the United States and efforts are underway to prevent establishment and spread (USDA 2019).

Invasions from species such as *V. mandarinia* are governed by arrival, establishment, and spread (Liebhold & Tobin 2008). Ecological niche models, which involve model calibration using climate variables in native ranges, followed by extrapolation to introduced areas, are often used to assess habitat suitability for invasive species (Leibhold & Tobin 2008; Peterson et al. 2013). Invasions can be particularly problematic in regions with high human activity, which can also facilitate invasions through transport of introduced species. To assess spread of invasive species, models can also simulate processes (Liebhold & Tobin 2008; Engler et al. 2012). Models thus guide efforts to prevent establishment and spread, which are cost-effective early in invasions (Liebhold & Tobin 2008).

It is not yet clear if *V. mandarinia* is established in North America, and federal and local agencies are implementing trapping and monitoring programs to identify areas of introduction and prevent establishment and spread (USDA 2019). However, several factors that could guide mitigation efforts remain unknown, including the potential habitat suitability for *V. mandarinia*. Moreover, the potential rate of population dispersal into new areas is poorly understood. In Europe, an invasion by the congener *V. velutina* has expanded via natural and human-assisted dispersal from 19 to nearly 80 km per year (Bertolino et al. 2016; Robinet et al. 2017). Here, we assess these questions by modeling responses of *V. mandarinia* to bioclimatic variables in the native range and extrapolating to introduced ranges. We also used dispersal simulations to estimate potential rates of spread throughout western North America. These complementary approaches can guide efforts to prevent the establishment and spread of this invasive species.

## Material and Methods

### Environmental factors affecting occurrence of V. mandarinia

We first assessed relationships between occurrence of *V. mandarinia* and environmental factors. Occurrence data were attained with the “spocc” package in R (R Core Team 2020) from the Global Biodiversity Information Facility, Biodiversity Information Serving Our Nation, Integrated Digitized Biocollections, and iNaturalist (Scott et al. 2017) (Fig. 1). Additional data were collected from published studies (Archer 1995; Lee 2010). Occurrence records located within oceans or without geographic coordinates were removed. 422 unique records from *V. mandarinia*’s native range in Asia (Japan, South Korea, Taiwan) were obtained (Fig. 1A). Of these, 200 were filtered out by enforcing a distance of 10 km between records (Fig. 1A); we used this filtering process because ecological niche models are sensitive to sample bias (Warren & Seifert 2011). Our assembled 222 records from east Asia are consistent with published records (Archer 1995; Lee 2010), suggesting we effectively captured the distribution of *V. mandarina*.

Vespine wasps have high endothermic capacity and thermoregulatory efficiency, and can survive broad temperature ranges (Käfer et al. 2012). To determine climate factors that constrain *V. mandarinia*, we obtained 7 Worldclim variables (Fick & Hijmans 2017): (i) annual mean temperature, (ii) mean diurnal range, (iii) max temperature of warmest month, (iv) minimum temperature of coldest month, (v) annual precipitation, (vi) precipitation of wettest and (viii) driest months (Bio14); we also considered annual mean radiation (Fig. S1A). Although some of these variables were correlated (Fig. S1B), highly correlated variables have little impact on ecological niche models that account for redundant variables (Feng et al. 2019).

After selecting variables, we used generalized linear models (GLM) with Bernoulli errors to model the probability of occurrence of *V. mandarinia* as a function of each bioclimatic factor. This approach was used to minimize the chances of overfitting models, and Hosmer Lemeshow goodness of fit test were used to evaluate GLM model performance (Hosmer et al. 1989). Rather than plotting a single partial response curve (i.e., fitting response curves for specific predictors while keeping the other predictors at their mean value), we adopted inflated response curves to explore species-environment relationships along the entire gradient while keeping the other predictors at their mean, minimum, median, maximum, and quartile values (Zurell et al. 2012).

### Realized niche modeling of native and introduced populations

After assessing environmental factors affecting *V. mandarinia* occurrence, we next assessed realized niches occupied by native and introduced populations. Given that only 4 occurrence points exist in the introduced range of western North America, two of which are within 10 km, we could not use a strict test of whether realized niches shifted during the introduction of *V. mandarinia* (i.e., niche equivalency and similarity test; Warren et al. 2010). Rather, we used minimum ellipsoid volumes to display and compare the two realized niches; this technique provides a clear vision of niche breadth for two populations and their relative positions in reduced dimensions (Qiao et al. 2016). We generated three environmental dimensions that summarized 90% of overall variations in the 8 global bioclimatic dimensions using principle component analysis in NicheA version 3.0 (Qiao et al. 2016).

### Ecological niche modeling

We used classical niche models to assess worldwide habitat suitability for potential spread of *V. mandarinia* (Peterson et al. 2013). Given that uncertainty exists with any individual model, we used an ensemble approach that averaged predictions of five algorithms: (i) generalized additive models, (ii) general boosted models, (iii) generalized linear models; (iv) random forests, and (v) maximum entropy models. Such ensemble models have been proposed as a consensus approach to more effectively estimate climate suitability, achieve a higher predictive capacity, and reduce uncertainty (Araújo & New 2007; Thuiller et al 2009; Zhu & Peterson 2017). To build models, 50% of observed records were used for model training and 50% for validation (Tsoar et al. 2007). We used a “random” method in *biomod2* to select 10,000 pseudo-absence records from “accessible” areas of *V. mandarinia* in Asia, which were delimited by buffering minimum convex polygons of observed points at 400 km (Owen et al. 2013). This selection of pseudo-absence records improves ecological niche model performance (Phillips & Dudík 2008).

For evaluation of models, we used Area Under the Curve (AUC) of Receiver Operating Characteristic (ROC) plots as a measure of model fit (Jiménez-Valverde et al. 2012). AUC has been criticized in niche model literature, and inference upon its values should be taken cautiously as we didn’t have reliable absence data (Leroy et al. 2012). However, here we simply tested niche model discriminability in native areas and not introduced areas. Final niche models were fitted using overall trimmed occurrence points for combination with footprint and displaying.

Habitat modification and disruption has been linked to invasiveness in some Vespidae, and invasions could be particularly problematic in regions with high human activity (Beggs et al. 2011, Robinet et al. 2017). In its native range, *V. mandarinia* is able to colonize green areas in cities, although at lower abundance than other vespine wasps (Choi et al. 2012). Human-assisted movement has affected expansion of *V. velutina* in Europe, and may also affect *V. mandarinia* (Robinet et al 2007). Models that combine climate suitability with measures of human activity may provide more accurate estimates of site vulnerability to colonization, particularly arrival and establishment processes (Liebhold & Tobin 2008). We measured human footprint as an indicator of human-mediated disturbances, a metric that combines population pressure and human infrastructure. We combined human footprint with climate suitability using a bivariate mapping approach (Fig. 2). All variables selected for analyses were used at a resolution of 5 arcmin.

**Figure 2.**
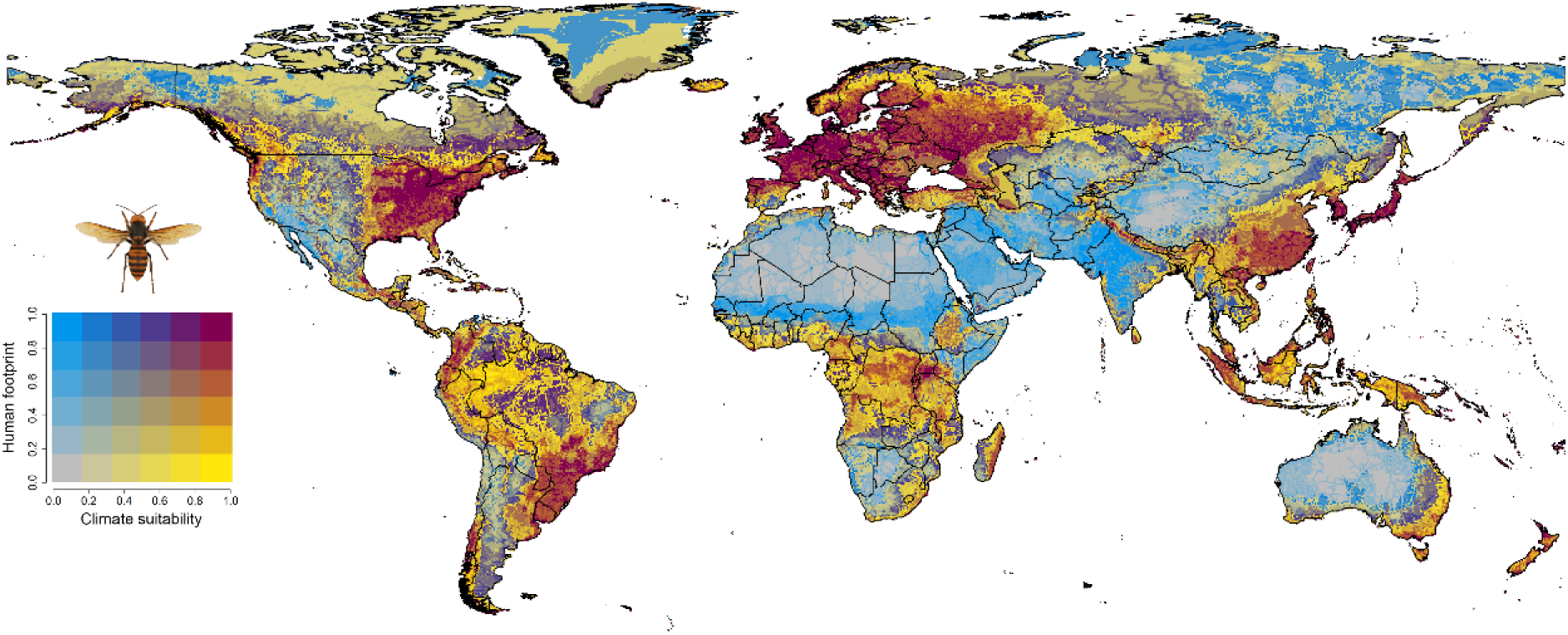
Ensemble forecast of potential invasion of *Vespa mandarinia*. Increasing intensities of yellow represent increasing climate suitability, and increasing blue represent increasing establishment potential due to human activity, where increasing red mean increasing potential invasion due to high climate suitability and human activity. Scores of bivariate maps are divided into 6 equal quantiles in the data ranges of climate suitability and human footprint respectively.

### Dispersal simulation

*Vespa mandarinia* is a social insect that forms colonies with one queen and many workers, and population dispersal is mediated by the spread of queens. Workers typically fly 1 to 2 km from their nest when foraging, although they can forage up to 8 km (Matsuura & Sakagami 1973). Data on queen dispersal appears to be unknown, but *V. mandarinia* queens are likely to have flight capacity greater than workers. Flight mill simulations with the congener *V. velutina* suggest that queens can fly 18 km in a single day (Robinet et al 2017), although flight distance under field conditions is likely to be smaller.

To simulate potential spread of *V. mandarinia* based on these dispersal capacities and occurrence points in western North America, we used the “MigClim” package (Engler et al. 2012). This approach simulates species expansion using a species’ occurrence as well as habitat suitability and different dispersal scenarios. Short-distance dispersal considers innate dispersal of a species to move through diffusion-based processes, whereas long-distance dispersal considers passive dispersal over long distance, such as dispersal by hitchhiking on human activity (Engler et al. 2012). MigClim uses a dispersal step as a basic time unit to simulate the dispersal, with dispersal steps often equal to one year since most organism dispersal occurs yearly or can be modeled as such, particularly for social insects where queens form colonies only once a year (Engler et al. 2012). We ran a simulation with a total of 20 dispersal steps for *V. mandarinia*.

In our simulations, we created combined suitability using climate suitability from ensemble models and human footprint. We then chose 3 different dispersal scenarios for simulations: (i) short-distance dispersal only, (ii) long-distance dispersal only, and (iii) both short- and long-distance dispersal. These three scenarios seek to capture both biological and human-mediated dispersal potential of *V. mandarinia*, as MigClim does not account for population demography (Engler et al. 2012). Simulations of short-distance dispersal were based on physical barriers and the dispersal kernel, which is the dispersal probability as a function of distance, whereas long-distance dispersal simulations depend on frequency of movement and distance range. MigClim uses a dispersal step as a basic time unit to simulate the dispersal, with dispersal steps often be equal to one year since most organism dispersal occurs yearly or can be modeled as such, particularly for social insects where queens form colonies only once a year (Engler et al. 2012).

We ran simulations with 20 dispersal steps. In MigClim, the dispersal kernel is the dispersal probability as a function of distance (*P*_*disp*)_ and the propagule production potential (*P*_*prop*_). Our raster data had a resolution of 5 arcmin (≈ 5.5 km); we defined short-distance dispersal as less than 6 pixels (∼ 33km). Dispersal more than 6 pixels was considered long-distance dispersal, which had a maximum 20 pixels (∼110km). We used a dispersal kernel of 1.0, 0.4, 0.16, 0.06, and 0.03 pixel for short-distance dispersal, which is an average of 10 km/dispersal step, with a maximum of 33 km. We set *P*_*prop*_ as 1 since *V. mandarinia* is a social insect and we assumed that the probability of a source cell to produce propagules is 100%. We assumed there were no barriers to either short- or long-distance dispersal.

## Results and Discussion

Generalized linear models showed no significant differences between models fit to the 8 environmental variables and observed data (χ^2^ = 8.2, df = 8, *P* = 0.41). We show *V. mandarinia* is most likely to occur in regions with low to warm annual mean temperatures and high annual precipitation (Fig. 2). However, our models show that they can tolerate broad temperature ranges (Fig. 2, S3), and that they are not particularly sensitive to radiation and extremes of precipitation (Fig. S3). The most suitable habitats are predicted to be in regions with maximum temperature of 39 °C in the warmest month (Fig. S3). Our results thus support the existence of a thermal threshold beyond which *V. mandarina* would be unable to establish, and this could be crucial for management and policy making in case of a prolonged invasion of the hornet in North America.

The minimum ellipsoid volumes show that the realized niche of introduced individuals in western North America were nested within the realized niche of native populations (Fig. 3). As the introduced locations represent a small fraction of the realized niche occupied by native populations, there is widespread potential for the introduced range to expand without mitigation (Fig. 3). However, the contrasting volume sizes occupied by native and introduced populations may simply be due to the limited number of occurrences outside of the native range (4 points) rather than any reduction in the niche space available to introduced populations.

**Figure 3.**
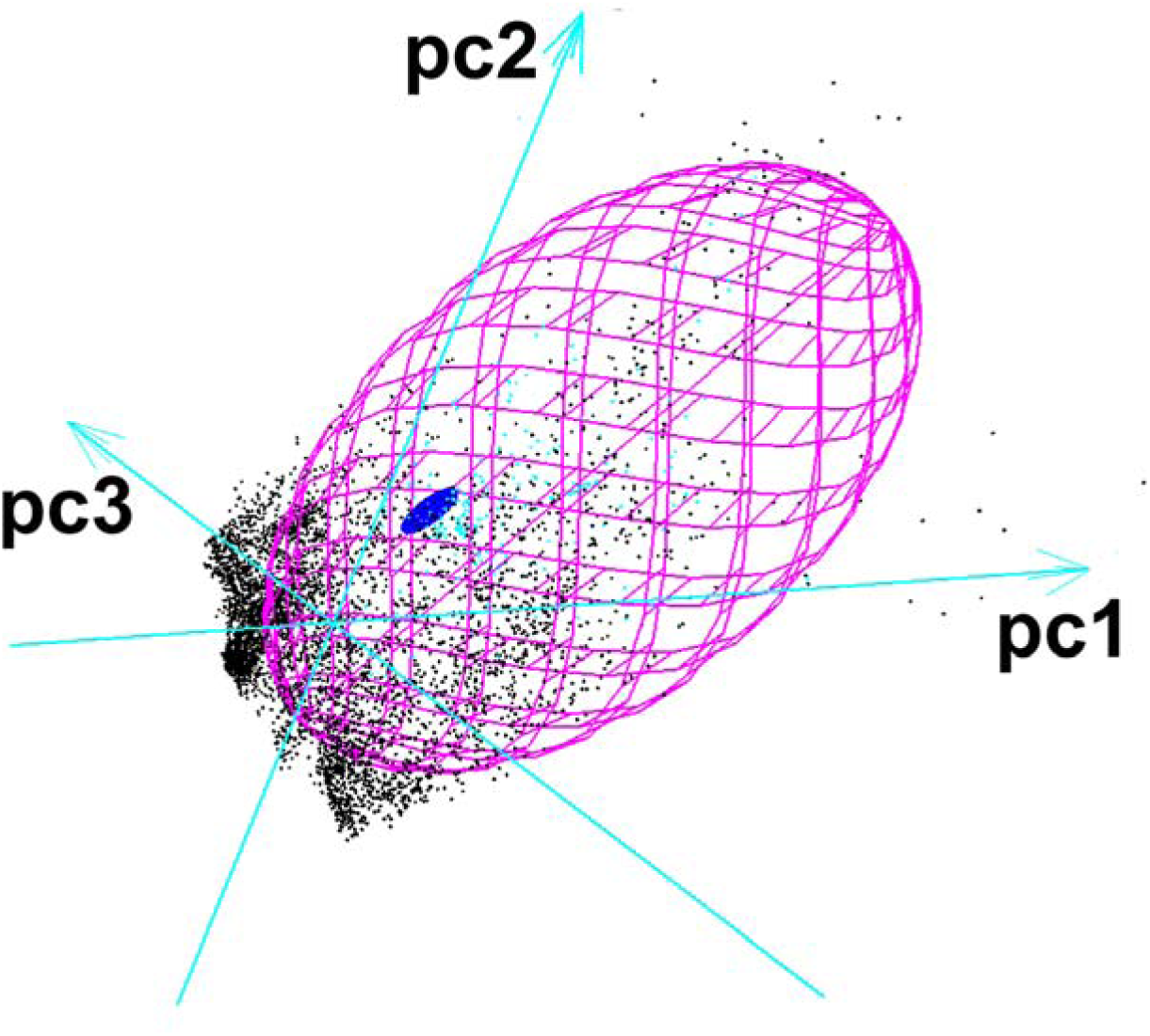
Realized niche occupied by native and introduced populations shown as minimum ellipsoid volumes. The pink volume represents the native niche, the blue volume represents the introduced niche, and points denote environmental conditions across the globe. The three PCA axes were estimated in NicheA and captured 90% of the variation in the 8 bioclimatic variables.

Our ecological niche models showed excellent performance in discriminability evaluations (generalized additive model [GAM]: AUC = 0.89; general boosted model [GBM]: AUC = 0.93; generalized linear model [GLM]: AUC = 0.91; Maxent: AUC = 0.93; Random Forest [RF]: AUC = 0.91). However, the five niche models had variability in habitat suitability across the globe (Fig. S4), and the ensemble model (Fig. 2) had better discriminability and outperformed these individual models (AUC = 0.94). Outside of the native area, our ensemble models captured detection points in North America as occurring in regions with highly suitable habitat (Fig. 2).

The ensemble models suggested that suitable habitat for *V. mandarinia* exists along much of the coastline of western North America as most of the eastern USA and adjacent parts of Canada, much of Europe, northwestern and southeastern South America, central Africa, eastern Australia, and most parts of New Zealand. Each of these regions is also associated with high human activity, although we did identify suitable climatic areas with low human activity (e.g., central South America, Fig. 2). Yet, given that many suitable regions were identified by the ensemble model that had both high climatic suitability and high human activity, it is likely that human activity could facilitate future invasions of *V. mandarinia*. The model predicts that much of the interior of North America is unsuitable habitat, likely due to inhospitable temperatures and low precipitation. This includes the eastern portions of British Columbia and the Pacific Northwest states, and the Central Valley of California, all of which have extensive agricultural production (e.g. tree fruit and tree nuts) that relies almost exclusively on *A. mellifera* pollination.

Our simulations of *V. mandarinia* dispersal in western North America showed high potential for spread within western North America (Fig. 4). When considering short-distance dispersal, mediated by hornets flying an average of 10 km/yr and a maximum of 33 km/yr, populations of *V. mandarinia* could rapidly spread along the western coast of North America, reaching Oregon in 20 yr. Northward expansion into Canada would likely be limited to the southern coast of British Columbia (Fig. 4). When we accounted for long-distance human-mediated dispersal, the expansion of *V. mandarinia* extended dramatically toward the north along coastal areas of British Columbia, and showed a faster rate of expansion into southern Washington State and into Oregon, USA (Fig. 4). This suggests dispersal throughout the western USA could occur within 20 or less yr even without human-mediated transport or new introduction events.

**Figure 4.**
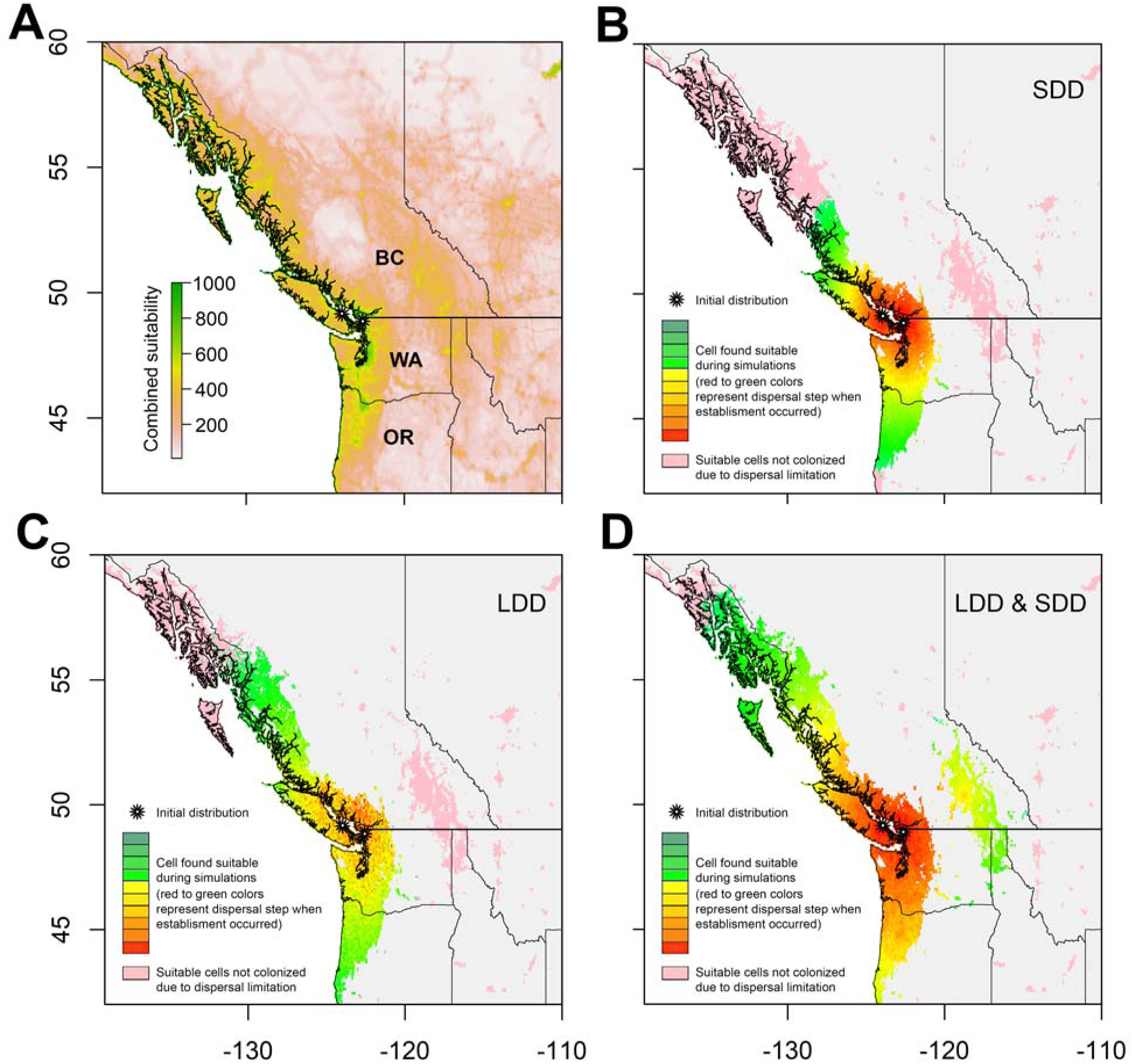
Combined suitability (A) and estimated expansion (B-D) of *Vespa mandarinia* over 20 yr in western North America under three dispersal scenarios: (B) short dispersal distance only (SSD), (C) long dispersal distance only (LDD) and (D) combined (LDD & SDD) scenarios. Each color represents two dispersal steps (total 20 dispersal steps) in dispersal simulations.

Ecological impacts are difficult to predict for vespids (Beggs et al. 2011). While many transplanted Vespidae appear to have only minor impacts, others are known to displace congeners through multiple, idiosyncratic mechanisms (Beggs et al 2011). There are no other *Vespa*, native or introduced, in the region of North America where *V. mandarinia* has been detected, and no native *Vespa* where suitable habitat is predicted by this model. However, Asian giant hornets are known to prey on social Hymenoptera other than bees (Matsuura 1984), and thus could affect populations of several vespid genera in the Pacific Northwest. *Vespa mandarinia* also preys upon many other insects, with chafer beetles comprising a large part of its diet in parts of Japan (Matsuura 1984). It is unknown how it might impact native insects if it becomes established, but the habitat suitability predicted here indicates that negative effects could be distributed over a fairly expansive area.

We also anticipate considerable impacts on beekeepers. Established populations of *V. mandarinia* would likely prey on readily-available hives late in the season, weakening any that aren’t killed outright. In Europe, the congener *V. velutina* has reportedly caused losses ranging from 18 to 80% of domestic hives, depending on the region (Laurino et al. 2020). The results presented here suggest that large expanses of the Pacific Coast in North America could become challenging for beekeeping operations, especially during the late summer and fall when attacks are greatest. Unchecked, this species of hornet could cause major disruption in the western US and Canada, possibly forcing beekeepers to invest in extensive hornet management or relocate parts of their operations to areas of unsuitable *V. mandarinia* habitat.

## Acknowledgement

We thank D. Zurell, R. Engler, and E. Ugene for help developing ecological models. The work was funded by USDA Hatch Project 1014754.

