## Supplementary Figures for "Assessing the ecological niche and invasion potential of the Asian giant hornet"

**Appendix S1**

**Figure S1.** Ranking of the 20 assembled bioclimate variables using the Boruta algorithm. The variables are (i) BIO1: annual mean temperature; (ii) BIO2: mean diurnal temperature range; (iii) BIO3- isothermality; (iv) BIO4: temperature seasonality; (v) BIO5: max temperature of warmest month; (vi) BIO6: min temperature of coldest month; (vii) BIO7: temperature annual range; (viii) BIO10: mean temperature of warmest quarter; (ix) BIO11: mean temperature of coldest quarter; (x) BIO12: annual precipitation; (xi) BIO13: precipitation of wettest month; (xii) BIO14: precipitation of driest month; (xiii) BIO15: precipitation seasonality; (xiv) BIO16: precipitation of wettest quarter; (xv) BIO17: precipitation of driest quarter; (xvi) Rmean: mean monthly radiation; (xvii) RMin: minimum monthly radiation; (xviii) Rmax: maximum monthly radiation; (xix) Rstd: standard deviation of monthly radiation; (xx) Rrange: range of monthly radiation.

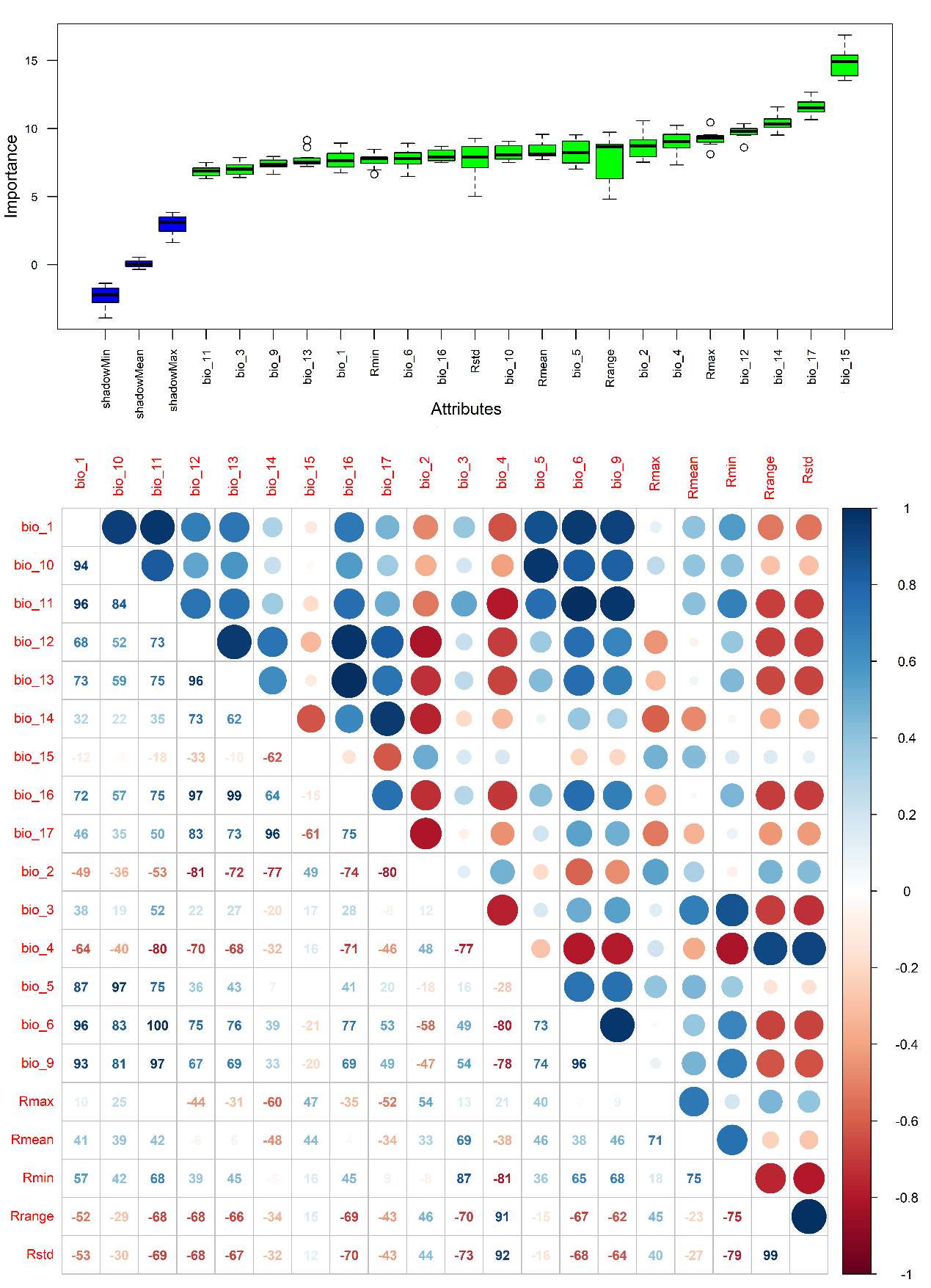

**Figure S2.** (A) Ranking of the 8 selected bioclimatic variables using the Boruta algorithm and (B) multi-collinearity among these 8 variables. The variables are (i) BIO1: annual mean temperature; (ii) BIO2: mean diurnal temperature range; (iii) BIO5: max temperature of warmest month; (iv) BIO6: min temperature of coldest month; (v) BIO12: annual precipitation; (vi) BIO13: precipitation of wettest month; (vii) BIO14: precipitation of driest month; (viii) rad: mean monthly radiation.

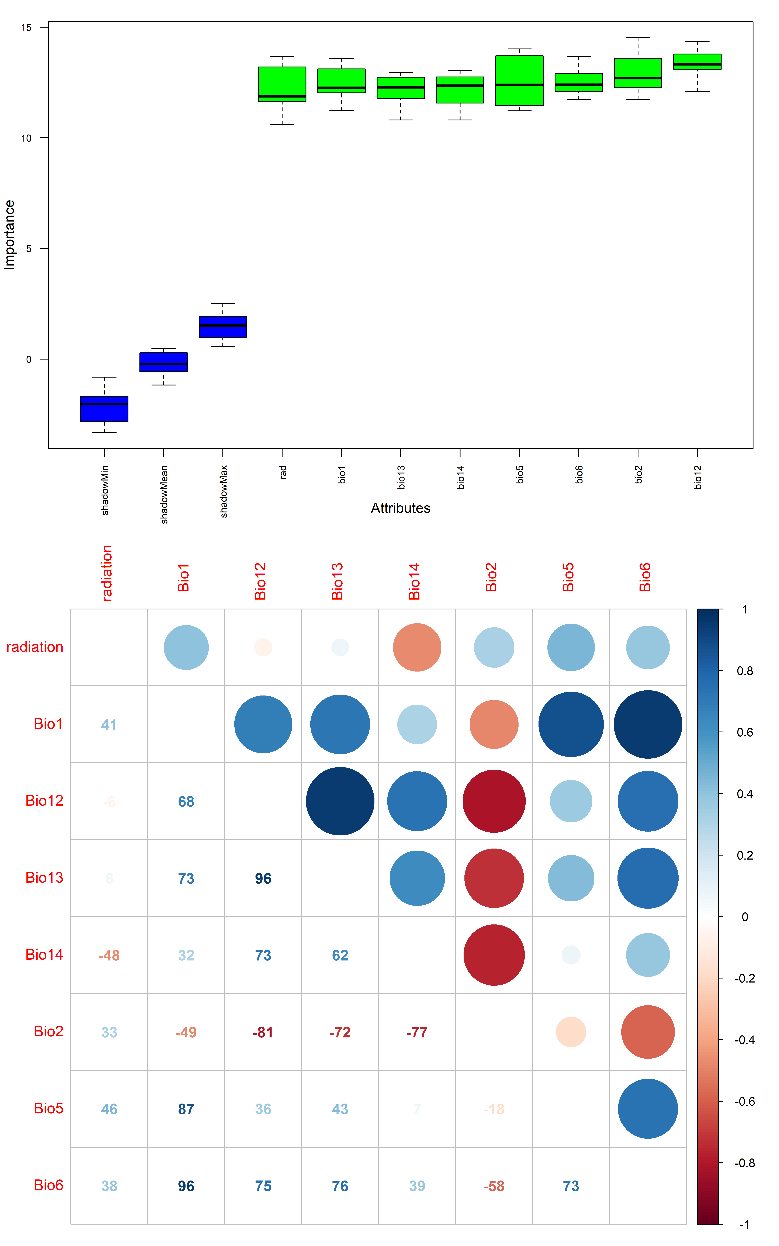

**Figure S3.** Inflated response curves of *Vespa mandarinia* to bioclimate variables. Blue lines denote partial response curves when holding other predictors at their mean value. Grey lines denote response curves when keeping other predictors at their minimum, median, maximum and quartiles values. Abbreviation of bioclimate refer to main text (units for Bio1, 2, 5, 6 are °C, for Bio12, 13, and 14 are mm, for radiation is kJ m^-2^ day^-1^).

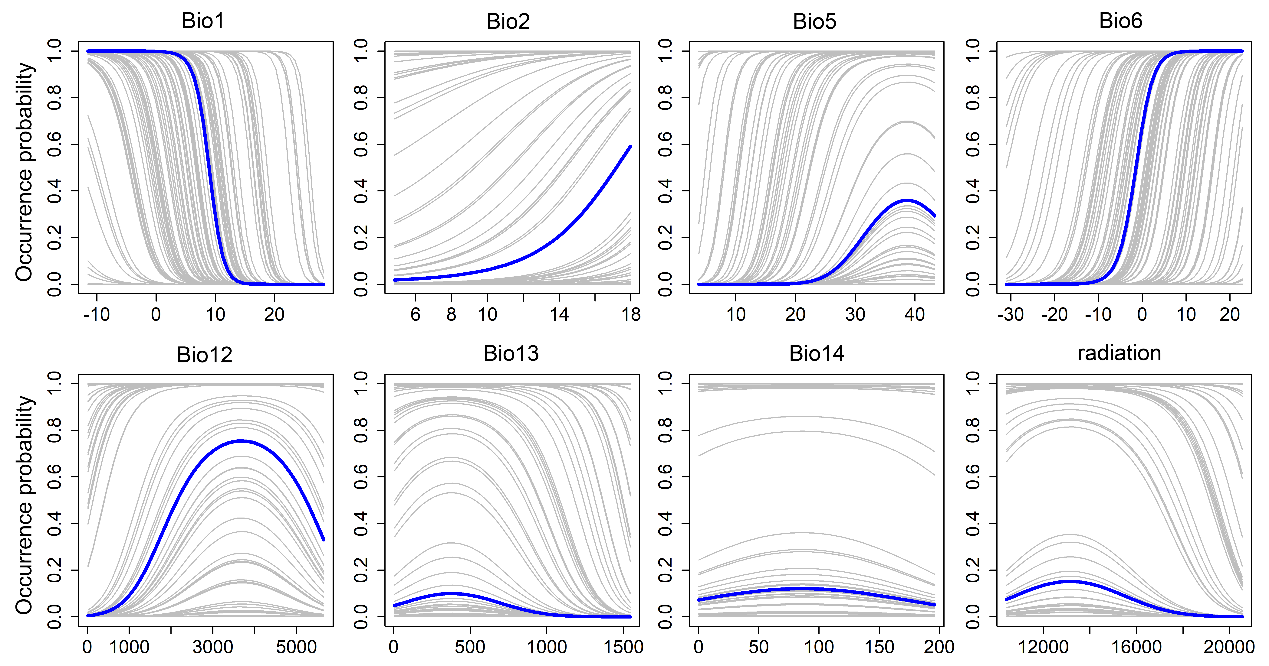

**Figure S4.** Spatial variations of niche model predications based 5 model algorithms. Warm color indicates high variation, cold color indicates low variation. Notice that the suitable areas identified by ensemble model.

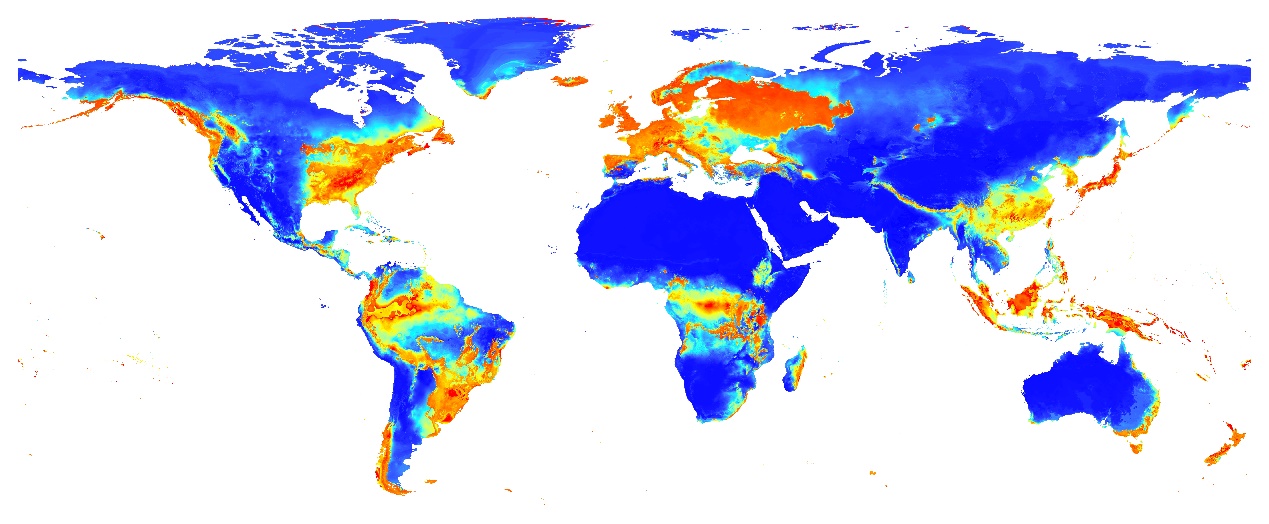
